# Zinc(II) binding on human wild-type ISCU and Met140 variants modulates Fe-S complex activity

**DOI:** 10.1101/262477

**Authors:** Nicholas G. Fox, Alain Martelli, Joseph F. Nabhan, Jay Janz, Oktawia Borkowska, Christine Bulawa, Wyatt W. Yue

**Affiliations:** Structural Genomics Consortium, Nuffield Department of Clinical Medicine, University of Oxford, UK OX3 7DQ; Pfizer Rare Disease Research Unit, Worldwide Research and Development, Pfizer Inc., 610 Main Street Cambridge, Massachusetts 02140

**Author notes:** Corresponding author: Wyatt Yue. Corresponding author: Christine Bulawa.

**Keywords:** Cysteine desulfurase, Friedreich’s ataxia, Iron-sulfur cluster, Zinc

## Abstract

The human *de novo* iron-sulfur (Fe-S) assembly complex consists of the cysteine desulfurase NFS1, accessory protein ISD11, scaffold protein ISCU, and allosteric activator frataxin (FXN). FXN has been shown to bind the NFS1-ISD11-ISCU complex (SDU), to activate the desulfurase activity and thus Fe-S cluster biosynthesis. Conversely, in the absence of FXN, the NFS1-ISD11 (SD) complex was reported to be inhibited by the binding of recombinant ISCU. Here, we show that recombinant ISCU binds zinc(II) ion, and that the presence of zinc in as-isolated ISCU has impacts on the SDU desulfurase activity as measured by sulfide production. Indeed, the removal of this zinc(II) ion from ISCU causes a moderate but significant increase in activity compared to SD alone, and FXN can activate both zinc-depleted and zinc-bound forms of ISCU complexed to SD. Recent yeast studies have reported a substitution on the yeast ISCU orthologue Isu, at position Met141 (Met140 in human numbering of precursor protein) to Ile, Leu, Val, or Cys that could bypass the requirement of FXN for Fe-S cluster assembly and cell viability. Using recombinant human proteins, we report no significant differences in the biochemical and biophysical properties observed between wild-type and variants M140I, M140 L, and M140 V of ISCU. Importantly, in the absence of FXN, ISCU variants behaved like wild-type and did not stimulate the desulfurase activity of the SD complex. This study therefore identifies an important regulatory role for ISCU-bound zinc in modulation of the human Fe-S assembly system in vitro but no ‘FXN bypass’ effect on mutations at position Met140 in human ISCU.

**ABBREVIATIONS:**

ACP: acyl carrier transfer protein
BLI: biolayer interferometry
BSA: bovine serum albumin
CD: circular dichroism
DMPD: NN-dimethyl-*p*-phenylenediamine
DSF: differential scanning fluorimetry
DTT: dithiothreitol; EDTA, ethylenediaminetetracetic acid
Fe-S: iron sulfur
FRDA: Friedreich’s ataxia
FXN: frataxin
HEPES: 4-(2-hydroxyethyl)-1-piperazineethanesulfonic acid
IPTG: isopropyl β-D-1-thiogalactopyranoside
PLP: pyridoxal 5′-phosphate
SD: protein complex composed of NFS 1 and ISD11
SDU: protein complex composed of NFS 1, ISD11
ISCU: SDUF, protein complex composed of NFS 1, ISD11, ISCU, and frataxin
TCA: trichloroacetic acid
TCEP: tris(2-carboxyethyl) phosphine
Tris: tris(hydroxymethyl)aminomethane

## INTRODUCTION

Iron-sulfur (Fe-S) clusters are prosthetic groups required for critical cellular functions including oxidative respiration, DNA repair, and the biosynthesis of other cofactors.^*1*^, ^*2*^ The Fe-S biosynthetic pathway in humans is located in the mitochondrial matrix and is initiated by a protein complex of NF<u>S</u>1, IS<u>D</u>11, ISC<u>U</u> and frataxin (<u>F</u>XN) (‘SDUF’).^*3*^–^*6*^ Recently, this protein complex was shown to be associated with an additional component, the acyl carrier protein (ACP, also known in human as NDUFAB1),^*7*^, ^*8*^ whether ACP is an intrinsic component of the complex *in vivo* remains to be determined. Within the complex, the cysteine desulfurase NFS1 (homolog of bacterial IscS or yeast Nfs1) catalyzes the pyridoxal phosphate (PLP)-dependent conversion of L-cysteine to L-alanine, and generates a persulfide species that delivers the sulfane sulfur to the Fe-S scaffold protein, ISCU (homolog of yeast Isu or bacterial IscU).^*9*^, ^*10*^ ISD11 (LYRM4) is a eukaryotic-specific protein belonging to the Leu-Tyr-Arg (LYR) superfamily of small and basic proteins, which interacts, stabilizes, and may regulate NFS1 activity.^*7*^, ^*11*^–^*16*^. Frataxin (FXN, homolog of yeast Yfh1 or *E. coli* CyaY) is an allosteric regulator of the Fe-S assembly complex, with human FXN shown to stimulate the rate of cysteine desulfurase activity^*17*^ and iron sulfur cluster biosynthesis^*18*^ *in vitro*, whereas the bacterial homolog CyaY appears to inhibit the counterpart desulfurase IscS.^*19*^

Within the de novo Fe-S assembly process, ISCU serves as the scaffold, obtaining ferrous iron and inorganic sulfide to assemble the Fe-S cluster. Human ISCU has three conserved cysteine residues namely Cys69, Cys95, and Cys138 (also referred as Cys35, Cys61, and Cys104 respectively in the literature^*^) involved in coordinating the Fe-S cluster. The LPPVK motif on ISCU is then recognized by the chaperone protein GRP75, to either deliver the Fe-S cluster to an apo-protein target, or possibly into the 4Fe-4S cluster pathway.^*20*^ A wealth of *E. coli* IscU and yeast Isu structures determined from NMR and X-ray crystallography studies^*21*^–^*30*^ have revealed two conformational states of the protein with respect to its secondary structures and local environment surrounding the conserved cysteines. The two conformations, namely structured and disordered, were observed with different liganded or mutant ISCU proteins, suggesting they may play a role in ISCU function and interaction.

Expansion of a GAA trinucleotide repeat in the *FXN* gene results in cellular depletion of frataxin protein and causes the autosomal recessive neurodegenerative disease Friedreich’s ataxia (FRDA). Recently, a “suppressor” mutation in *S. cerevisiae* Isu, corresponding to the substitution of Met141 (also referred as Met107 in the literature^*^) to either an Ile, Leu, Val, or Cys, was shown to increase cell viability and rescue Fe-S cluster synthesis in Yfh1 (FXN equivalent)-deleted cells.^*31*^–^*33*^ Phylogenetic analysis of ISCU sequences showed that in prokaryotes, the equivalent position to yeast Met141 is more commonly an Ile, Leu, Val or Cys. At the amino acid sequence level, substitution of Met141 with the aforementioned residues yielded the yeast suppressor variant of Isu that behaved like the prokaryotic orthologue and facilitated cluster assembly without frataxin.^*32*^ Transfer of the persulfide from NFS1 to the proposed sulfur acceptor (i.e. ISCU Cys138) ^*34*^, ^*35*^ is positioned only two residues away from ISCU Met141 and a bacterial structure of IscU has shown to adopt a conformation change around this region when a [2Fe-2S] cluster is bound.^*26*^ A substitution of the nearby Met residue could therefore impose a direct effect on this crucial functional interaction.

The objectives of this study are two-fold, first to investigate the biochemical and biophysical properties of recombinant human ISCU towards its functional interaction within the SDUF complex for Fe-S cluster assembly, and second to determine if the recombinant Met141 substitution variants of human ISCU result in frataxin bypass *in vitro*. We also report a discovery of a role for zinc(II), co-purified with recombinant ISCU wild-type and variant proteins, on NFS1 desulfurase activity.

## EXPERIMENTAL PROCEDURES

### Cloning, expression and purification of human ISCU, FXN, and NFS1-ISD11

Constructs of human ISCU (Δ1-34) and Frataxin (Δ1-80) were subcloned into the pNIC28-Bsa4 vector (GenBank ID: EF198106) for recombinant *E. coli* expression using BL21(DE3)-R3-pRARE2 cells. For bi-cistronic co-expression of NFS1-ISD11, a DNA fragment encoding His-tagged ISD11 and non-tagged NFS1 (Δ1-55), separated by an in-frame ribosomal binding site, was sub-cloned into pNIC28-Bsa4 vector. ISCU variants (M140I, M140 L, and M140 V) were constructed using the QuikChange site-directed mutagenesis kit (Strategene), and confirmed by DNA sequencing. Cells transformed with the above plasmids were grown in Terrific Broth and induced with 0.1 mM isopropyl β-D-1-thiogalactopyranoside (IPTG) for 16 hours at 18 °C. Cell pellets were resuspended in binding buffer (50 mM HEPES pH 7.5, 500 mM NaCl, 20 mM Imidazole, 5% glycerol, and 2 mM TCEP) containing EDTA-free protease inhibitor (Merck), and lysed by sonication. For NFS1-ISD11 purification, 150 μM pyridoxal 5’-phosphate (PLP) was supplemented to the binding buffer during sonication. The clarified supernatant was incubated with 2.5 mL Ni Sepharose 6 fast flow resin (GE Healthcare), washed and eluted with binding buffer containing 40 mM and 250 mM Imidazole, respectively. Elution fractions were collected, 10 mM DTT was added to samples containing ISCU or NFS1, and loaded onto gel filtration (Superdex S75 for ISCU and FXN, Superdex S200 for NFS1-ISD11 complex; GE Healthcare). Peak fractions were collected, treated with His-TEV protease to remove His-tag, and then passed onto Ni Sepharose 6 fast flow resin to remove His-TEV and cleaved His-tag. Fractions containing target protein were collected and buffer exchanged into gel filtration buffer (50 mM HEPES pH 7.5, 200 mM NaCl, 5% glycerol, and 2 mM TCEP).

### Methylene blue activity assay

Sulfide production, due to cysteine desulfurase enzyme activity, was measured using the methylene blue colorimetric assay as described previously.^*3*^, ^*36*^ The assay was run in buffer consisting of 50 mM HEPES pH 7.5, 200 mM NaCl, 10 mM DTT and either 100 μM EDTA or 50 μM ZnCl_2_. When noted, concentration of NFS1-ISD11 (SD) was at 0.5 μM, ISCU (U) at 2.5 μM, and Frataxin (F) at 40 μM was mixed in a 1.5 mL black Eppendorf tube with total volume of 800 μL. To allow comparison, the same excess of ISCU (5 equivalents to [SD]) and FXN (80 equivalents to [SD]) were used for all wild-type and mutant ISCU. The reaction was initiated by adding 100 μM L-Cysteine and placed in 37 °C incubator for 10 min (with FXN) or 20 min (without FXN) and then quenched with 100 μL of 30 mM FeCl_3_ in 1.2 N HCl and 100 μL of 20 mM *N,N*-dimethyl-*p*-phenylenediamine (DMPD) in 7.2 N HCl and placed back in 37 °C incubator for 20 min, followed by centrifugation to spin down precipitant and then take the absorbance at 670 nm. Concentration of sulfide was calculated *via* a standard curve of Na_2_S. Concentrations for ISCU and FXN were determined by a titration until maximum flux in activity was seen.

### SDU complex reconstitution by size exclusion chromatography

Reconstitution of the recombinant NFS 1-ISD11-ISCU complex (SDU) was mediated by co-expression of all three proteins in a poly-cistronic fashion, where only ISD 11 is His-tagged. Expression and affinity chromatography were carried out as described above for single proteins. Complex-containing fractions eluted from affinity step were pooled, and analytical gel filtration was performed using a Dionex Ultimate ™ 3000 system. The Sepax SRT SEC-300 7.8x300 mm column was pre-equilibrated in buffer containing 50 mM HEPES pH 7.5, 150 mM NaCl, 5% glycerol and 2 mM TCEP, and run at 0.5 mL/min. Complex formation was confirmed by TCA-precipitation followed by SDS-PAGE analysis.

### Differential scanning fluorimetry

Miniaturized DSF (nanoDSF) was performed in 10 μL capillaries using the Prometheus device (NanoTemper Technologies) that uses excitation at 280 nm to measure emission from tryptophan and tyrosine residues at two wavelengths: 330 (non-polar environment emission) and 350 nm (polar environment emission). Each capillary consists of ISCU protein at 0.8 mg/mL in buffer containing 50 mM HEPES pH 7.5, 200 mM NaCl and 10 mM DTT, supplemented with either 300 μM EDTA or 100 μM ZnCl_2_. Thermal unfolding was carried out using a linear thermal ramp (1.0 °C/min; 20 °C to 95 °C) and unfolding midpoint (melting temperature, T_m_) was determined from changes in the emission wavelengths of tryptophan and tyrosine fluorescence at 330 and 350 nm.

### Circular Dichroism

Circular dichroism (CD) spectra were recorded on a J-815 spectropolarimeter (JASCO) at 20°C with a scan speed of 100 nm/min using 0.1 cm pathlength quartz cells from Starna Scientific UK. The concentration of ISCU protein was at 0.1 mg/mL in buffer which consisted of 50 mM HEPES pH 7.5, 200 mM NaCl, 5 mM DTT and either 100 μM EDTA or 30 μM ZnCl_2_ and then buffer exchanged into 10 mM Tris pH 7.5 (the pH was adjusted with phosphoric acid) and 50 mM NaF before acquiring data. Data points were collected with a resolution of 0.2 nm, an integration time of 1 second, and a slit width of 1 nm. Each spectrum shown is the result of nine averaged consecutive scans, from which buffer scans were subtracted.

### Biolayer Interferometry (BLI)

BLI experiments were performed on a 16-channel ForteBio Octet RED384 instrument at 25 °C, in buffer containing 50 mM HEPES pH 7.5, 200 mM NaCl, 5% Glycerol, 2 mM TCEP, 5 mM DTT, 0.5 mg/mL BSA, which is further supplemented with either 100 μM ZnCl_2_ or 300 μM EDTA. 50 μL of 1 mg/mL of biotinylated ISCU was diluted to 650 μL and loaded to the streptavin coated sensors. The concentration for SD used ranged from 10 μM to 1.56 nM. Measurements were performed using a 300 second association step followed by a 300 second dissociation step on a black 384-well plate with tilted bottom (ForteBio). The baseline was stabilized for 30 sec prior to association and signal from the reference sensors was subtracted. A plot of response vs. [SD] was used for K_d_ determination using one site-specific binding fit in GraphPad Prism (GraphPad Software).

## RESULTS

### Recombinant ISCU is purified as a mixture of zinc-depleted and zinc-bound forms

*E. coli* IscU was previously shown to exist in two interconvertible conformations, comprising a structured and a disordered form that can be influenced by the zinc loading status.^*21*^, ^*24*^, ^*25*^ We set out to investigate if similar zinc-binding properties exist for human ISCU, a 167-aa precursor protein (133 aa mature protein without the mitochondrial signal peptide) bearing 75% sequence identity to *E. coli* IscU (Supplemental Figure 1). Analysis of as-purified ISCU by native mass spectrometry showed that the recombinant protein exists in two forms, one corresponding to the expected mass of the apo-protein (14604 Da), and the other with an additional mass of 64 Da (MW = 14668 Da) that suggests a zinc-bound form with one Zn^2+^ per ISCU monomer (Figure 1A top). The presence of the metal was further confirmed by treatment of the recombinant protein with EDTA or supplementation with ZnCl_2_ and analysis by native mass spectrometry (Figure 1A middle and bottom, respectively). Nano differential scanning fluorimetry demonstrated that the zinc-bound form of ISCU is significantly more thermostable with a melting temperature (T_m_ of 65.7 ± 0.2 °C) ~30 °C higher than that for the zinc-depleted form (T_m_ of 33.2 ± 0.6 °C) (Figure 1B,D). The metal-conferred thermostability was also reflected by far-UV circular dichroism spectra consistent with zinc-bound ISCU displaying a higher helical percentage (28% compared to 20%) and less disordered percentage (35% compared to 41%) than the zinc-depleted ISCU (Figure 1 C,D). There was no significant change in the percentage of β-sheets, suggesting that the observed changes in secondary structure are located around the Fe-S cluster binding site, consistent with previously reported bacterial structures of IscU with and without zinc.^*7*^, ^*23*^

**Figure 1:**
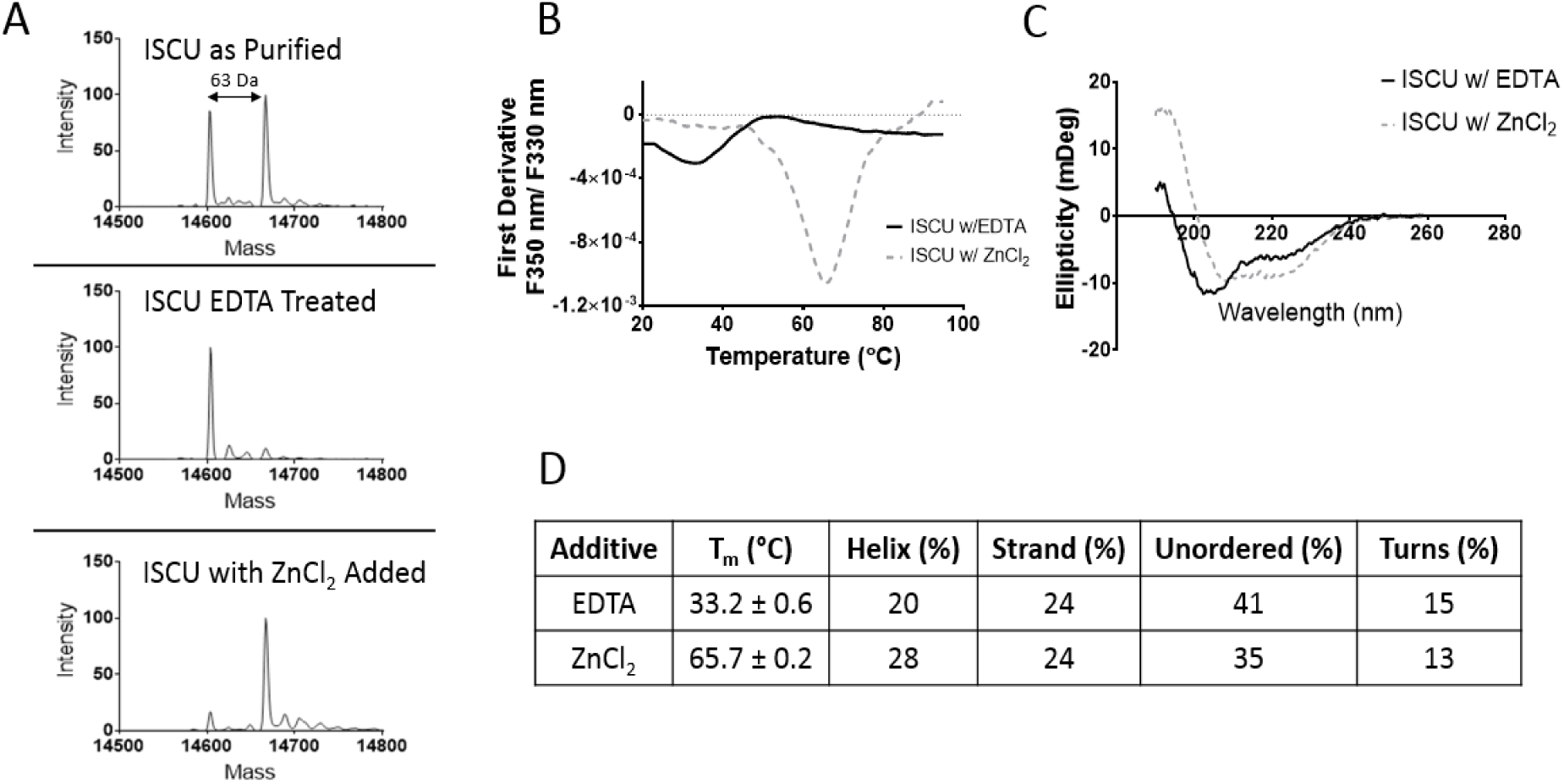
Characterization of wild-type ISCU in zinc-depleted and zinc-bound forms. (A) Mass Spectrometry analysis of purified ISCU shows the existence of two states with a mass difference of 64 Da consistent with a zinc(II) ion being bound. EDTA can successfully remove the zinc, and on the contrast the addition of ZnCl_2_ can push it to the zinc-bound state. (B) To determine thermostability of the two states, nanoDSF was used to determine the melting temperatures showing that zinc-bound was ~30 °C higher than that for the zinc-depleted form (T_m_ of 65.7 ± 0.2 °C, 33.2 ± 0.6 °C, respectively) and thus much more stable. (C) Circular Dichroism was used to determine secondary structure of the two states and found that the zinc-bound state had a higher helical percentage and a lower disordered percentage than that of the zinc-depleted state. (D) The table to outline the values obtained of the analysis of data from Figures 1 B&C.

### ISCU modulates NFS1-ISD11 activity *in vitro* in a zinc-dependent manner

In the first steps of Fe-S cluster assembly, ISCU functions as a scaffold protein for the NFS1-ISD11 desulfurase complex which generates and transfers the inorganic sulfur to ISCU, in a reaction that is activated by FXN protein.^*3*^, ^*37*^ We tested if the presence of zinc(II) ion in purified samples influences the desulfurase activity of NFS1-ISD11 using a methylene blue activity assay that measures sulfide formation,^*3*^, ^*36*^, ^*38*^, ^*39*^ in reaction buffer supplemented with either EDTA (100 µM) or ZnCl_2_ (50 µM) (Figure 2A). Recombinant human NFS1-ISD11 co-purified with bacterial acyl-carrier protein (ACP) from the host expression organism *E. coli* (Supplementary Figure 5), consistent with previous reports.^*7*^, ^*8*^ It is of note that the association with bacterial ACP has not been shown to influence NFS1 desulfurase activity. The NFS1-ISD11 complex (SD) displayed a basal level of desulfurase activity in both buffers (1.23 ± 0.08 mol/SD/min in EDTA-containing buffer; 1.64 ± 0.17 mol/SD/min in ZnCl_2_-containing buffer). The addition of ISCU to NFS1-ISD11 (SDU) increased the desulfurase activity by ~1.5 fold when assayed in EDTA-containing buffer. To our surprise, when SDU was assayed in ZnCl_2_-containing buffer, desulfurase activity was almost completely abolished. The activity of SD alone, without ISCU, was not affected by increasing ZnCl_2_ concentration (Figure 2A and Supplemental Figure 2), indicating that the zinc effect is mediated directly through the ISCU protein, and not due to indirect solution properties (e.g. metal-induced aggregation of the SD sample).^*40*^ As expected, further addition of the activator protein FXN (at 80 equivalents) to SDU (‘SDUF’) increased desulfurase activity in both buffer conditions, reaching a maximal activity of 8.14 ± 0.42 mol/SD/min in EDTA-containing buffer (4.5-fold increase from SDU) and 6.42 ± 0.16 mol/SD/min in ZnCl_2_-containing buffer (30-fold increase from SDU) (Figure 2A,C and Table 1).

**Table 1:**
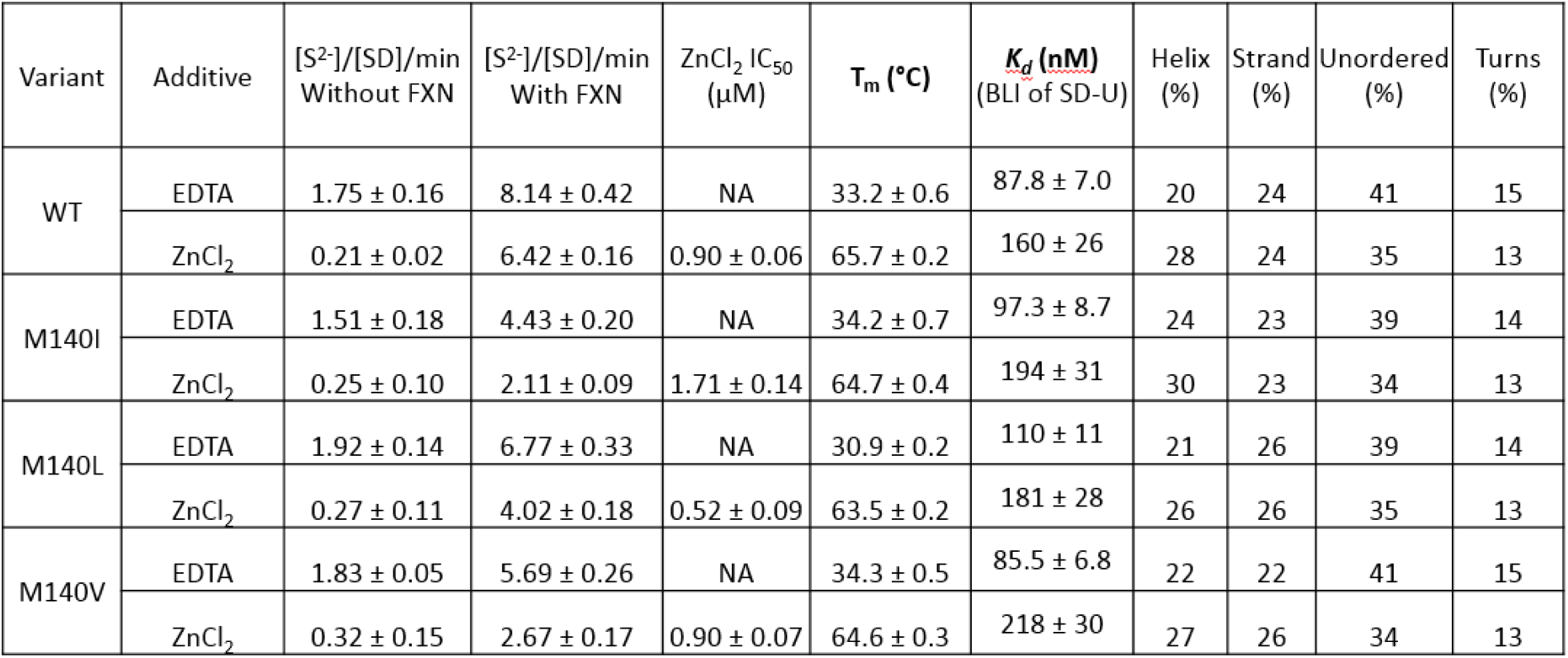
A Comparison on the Biophysical Properties of WT-ISCU and Proposed Suppressor Variants

Our result indicates that the inhibition of SD desulfurase activity by ISCU *in vitro*, an observation previously reported by various groups,^*3*^, ^*19*^, ^*37*^, ^*41*^ is a function of the divalent zinc(II) ion. To quantify zinc-mediated inhibition, ISCU was pre-incubated with EDTA, buffer exchanged, and assayed for desulfurase activity in buffer with increasing ZnCl_2_ concentrations (Figure 2B). Zinc(II) ion demonstrated a dose-dependent inhibition on desulfurase activity with an IC50 of 0.90 μM ± 0.06 for ZnCl_2_ (Figure 2B). To determine if the inhibition of SD activity was due to disrupted protein-protein interactions between ISCU and the NFS1-ISD11 complex, we characterized the binding between SD and biotinylated ISCU using bio-layer interferometry (BLI) (Figure 2 C, Supplemental Figure 3A). We did not observe any significant difference between the two forms of ISCU in their binding to SD, with dissociation constants in the nanomolar range: 87.8 ± 7.0 nM in buffer containing EDTA and 160 ± 26 nM in buffer containing ZnCl_2_.These findings suggest that complex formation was not affected in the assay at the tested Zinc(II) and protein concentrations.

**Figure 2:**
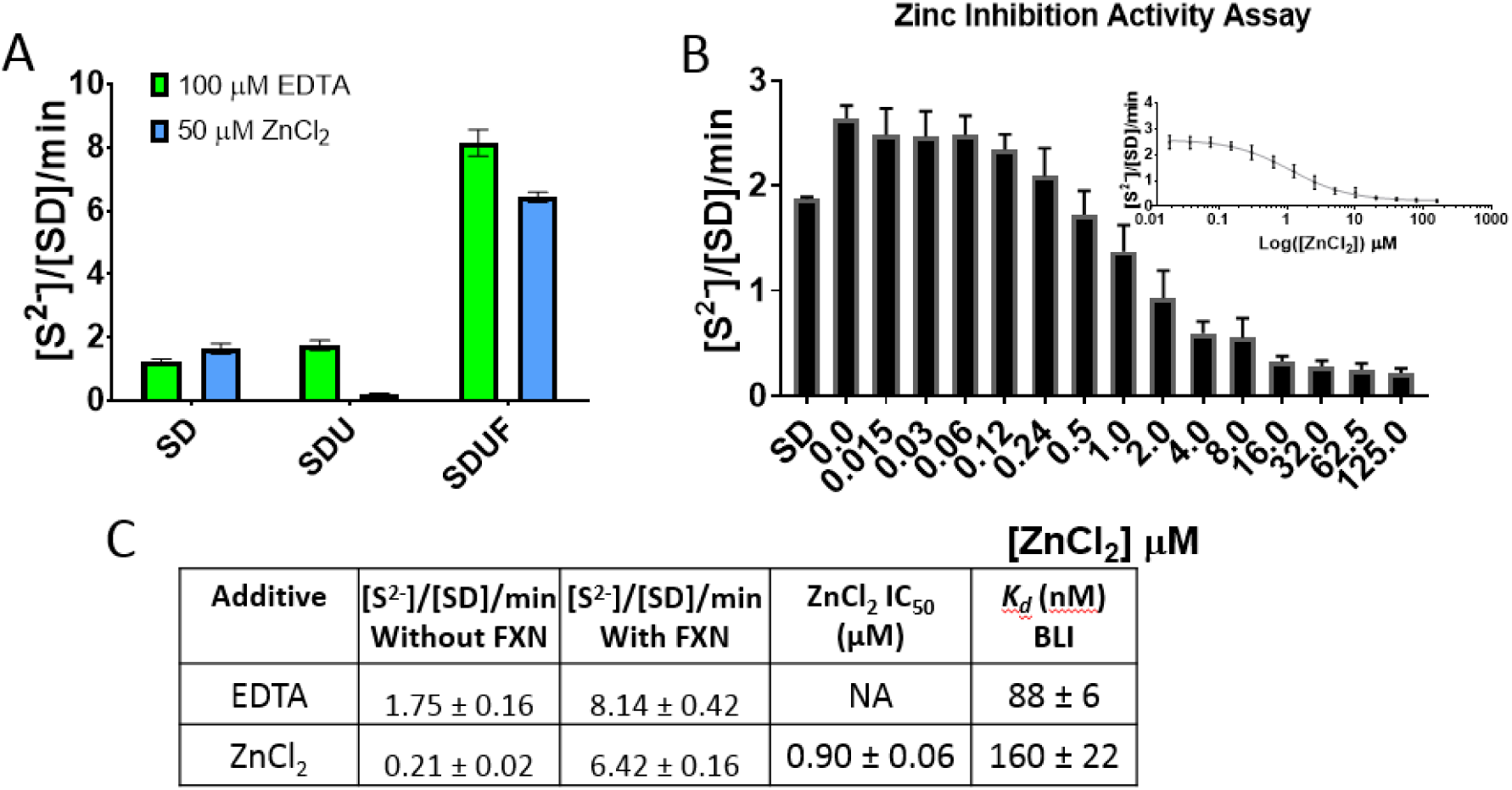
Activity assay analysis of wild-type ISCU in zinc-depleted and zinc-bound forms. (A) The methylene blue assay was used to determine the effect of desulfurase activity on the two states of ISCU using buffer supplemented with either 100 μM EDTA or 50 μM ZnCl_2_. When noted, concentration of NFS 1-ISD11 (SD) was at 0.5 μM, ISCU (U) at 2.5 μM, and Frataxin (F) at 40 μM. (B) The methylene blue assay was used to determine IC_50_ values for zinc to the NFS 1-ISD11-ISCU (SDU) by using buffer containing a serial dilution of zinc concentrations and plotting [inhibitor] *vs.* response (inset shown on a log scale). (C) The table to show values obtained from Figures 2A&B and binding constants obtained from BLI shown in Supplemental Figures 3.

### Human ISCU variants M140I, M140 L and M140 V behave as wild-type *in vitro*

In frataxin-deleted yeast, a substitution on the scaffold protein IscU at position Met 141 to either Cys, Ile, Leu, or Val, corrected the loss-of-frataxin phenotypes and rescued cell viability.^*32*^, ^*33*^ The IscU suppressor variant protein was also shown to bind and activate Nfs1 *in vitro* using purified yeast proteins, hence bypassing the requirement of Yfh1.^*42*^ In the human ISCU amino acid sequence (bearing 72% identity to yeast), the equivalent residue is Met140 (previously referred to as Met106; Supplemental Figure 1). Human ISCU variants with the corresponding Met substitutions (ISCUm_140_i, ISCUm_140_ l and ISCUm_140_ v) were constructed and expressed in *E. coli* as for wild-type (ISCUwt). All ISCU variants were purified as a mixed population of interconvertible zinc-depleted and zinc-bound forms (Supplemental Figure 4A,B), which behaved like ISCUwt in terms of secondary structure properties (Figure 3A, Table 1) and zinc-mediated thermostability (Table 1). All ISCU variants also retained the ability to form a stable complex with SD in size exclusion chromatography (Figure 3B and Supplemental Figure 5). Altogether, recombinant ISCU wildtype and variant proteins have indistinguishable structural and biophysical properties.

**Figure 3:**
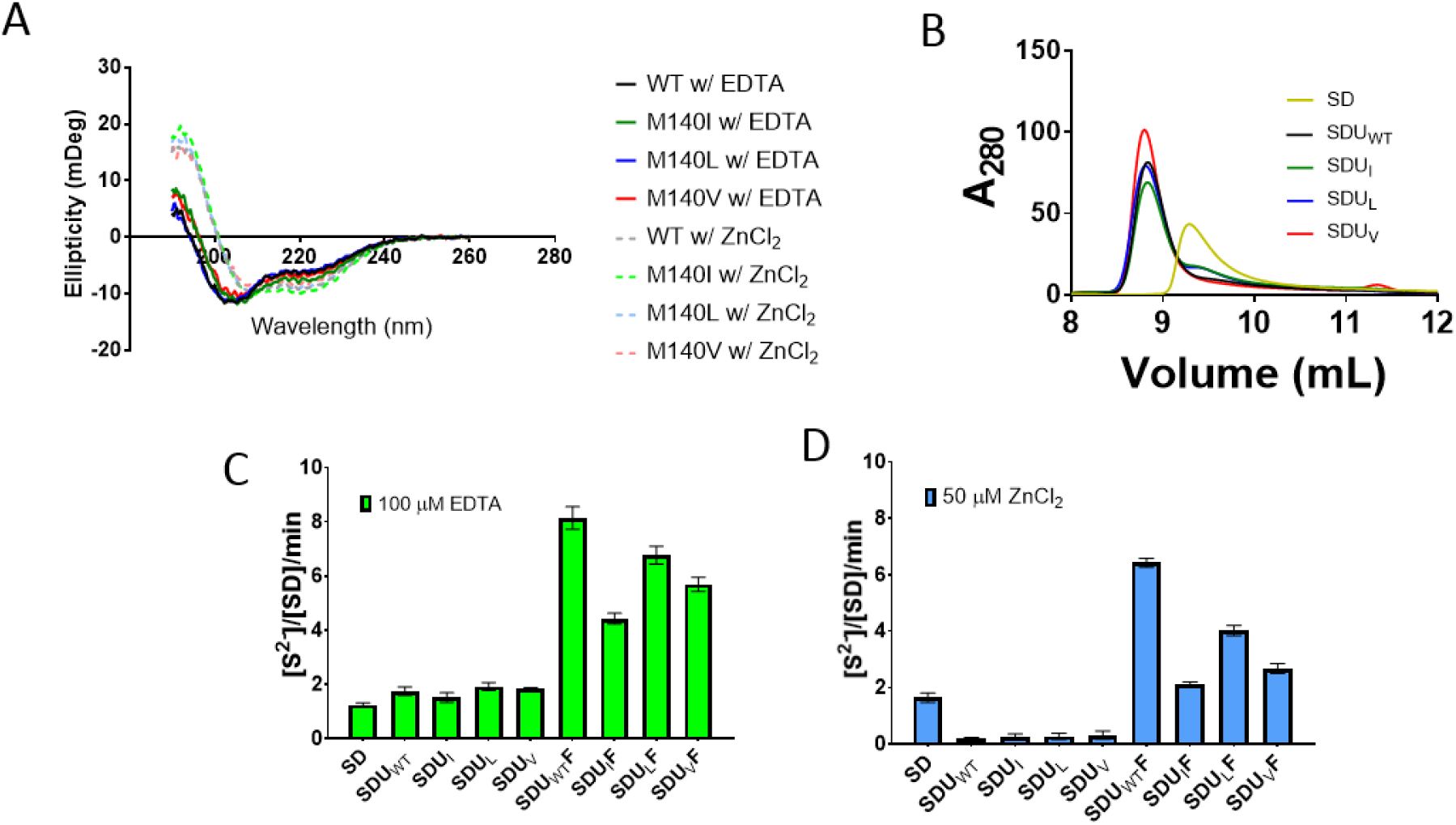
A biophysical approach to compare WT-ISCU to the human equivalent of the yeast-surpressor variants. (A) Circular Dichroism was used to determine secondary structure of the two states of ISCU and compare the WT to the M140I, M140 L, and M140 V variants and found no difference between them and showed similarly that the zinc-bound state had a higher helical percentage and a lower disordered percentage than that of the zinc-depleted state. (B) Analytical gel filtration was used to determine if complex formation was intact for NFS-ISD11 (SD), NFS1-ISD11-ISCUwt (SDUwt), NFS1-ISD11-ISCUm140i (SDUi), NFS1-ISD11-ISCUm_140_ l (SDUl), and NFS1-ISD11-ISCU_M_14_0 V_ (SDUV). (C,D) The methylene blue assay was used to determine the effect of desulfurase activity on the two states of ISCU WT and variants using buffer supplemented with either 100 μM EDTA (C) or 50 μM ZnCl_2_ (D). When noted, concentration of NFS1-ISD11 (SD) was at 0.5 μM, ISCU (U) at 2.5 μM, and Frataxin (F) at 40 μM.

We next employed the methylene blue activity assay to determine the effect of ISCU variants on the SD complex activity. All ISCU variants behaved similar to wild-type, in their respective zinc-depleted forms, and compared to SD-alone, increased activity slightly by ~1.5 fold (Figure 3 C). Importantly, in their zinc-bound forms SD activity was once again abolished (Figure 3D, Table 1). Therefore, the ISCU variants inhibited SD activity in a zinc-dependent manner, with IC_50_ values for ZnCl_2_ within the same order of magnitude as observed for SD in the presence of wild-type ISCU (Supplemental Figure 6, Table 1). The SDU complexes containing ISCU variants retained the ability to be activated by frataxin under both buffer conditions tested, with the degree of activation (evaluated by the maximal desulfurase activity reached in each case) in the following decreasing order: ISCU_WT_ (i.e. SDU_WT_F complex) > ISCU_M140 L_ (i.e. SDU_L_F complex) > ISCU_M140 V_ (i.e. SDU_V_F complex) > ISCU_M140I_ (i.e. SDU_I_F complex) (Figure 3 C and 3D, Table 1).

## DISCUSSION

In this study we set out to characterize recombinant human ISCU wild-type and variant proteins for their functional interaction with NFS1-ISD11 and effect on NFS1 activity. Our biophysical characterization of human ISCU is consistent with the existence of two conformational states, one being structured and one disordered around the Fe-S cluster binding site.^*43*^ Several structures of bacterial IscU also reveal a more structured conformation such as when having an Fe-S cluster bound (PDB 2Z7E),^*26*^ in complex with IscS (PDB 3LVM),^*27*^, ^*28*^ carrying a substitution on Asp39 (*E.coli* numbering, human equivalent Asp71; PDB 2KQK),^*23*^ or most commonly when bound to zinc(II) ion in place of an Fe-S cluster *via* the conserved cysteine residues (e.g. PDB 1WFZ, 1RP9P, 1XJS, 1SU0).^*29*^, ^*30*^ The more flexible/disordered conformation is observed in an NMR structure of wild type *E. coli* apo-IscU (PDB 2L4X) undergoing a helix to coil rearrangement in the helix containing the sulfur acceptor residue Cys106 (*E.coli* numbering, human equivalent Cys138).^*23*^

During the early steps of Fe-S cluster assembly, the cysteine residues on ISCU are responsible for accepting the persulfide product generated from NFS1 desulfurase, to be either incorporated in an Fe-S cluster, or reductively cleaved to release the sulfide ion. In the eukaryotic process, frataxin is known to activate the cysteine desulfurase activity of NFS1, thus up-regulating Fe-S cluster assembly.^*3*^, ^*34*^ Previous publications have shown that ISCU, in the absence of frataxin, inhibits the NFS1 desulfurase activity *in vitro*.^*34*^, ^*37*^, ^*44*^ Our study here attributes this inhibitory property of ISCU to the presence of zinc(II) ions bound to ISCU protein from recombinant expression, likely associated with the active site cysteine residues as observed in the several crystal structures of zinc-bound bacterial IscU. One possible explanation is that the zinc(II) ion bound to ISCU cysteine residues would prevent the sulfur acceptor residue Cys138^*34*^ from reductively cleaving the persulfide off NFS1, resulting in NFS1 inhibition and lack of catalytic turnover. Our data suggest no significant difference between the two ISCU forms in their binding affinity to the SD complex, nor any effect of zinc(II) on complex activity. Altogether, our findings suggest that zinc-bound ISCU directly inhibits NFS1 cysteine desulfurase activity.^*7*^ Alternatively, the bound zinc(II) ion could inhibit NFS1 activity by locking its mobile loop active site cysteine. Consistent with this, the recently published crystal structures of NFS1-ISD11-ACP-ISCU, in the presence and absence of zinc,^*7*^ reveal the zinc(II) ligation to include three ISCU residues (Cys95, His137, and Asp71, human precursor numbering) as well as the mobile loop cysteine from NFS1.^*7*^ This would imply that the mobile loop cysteine on NFS1 is sequestered by zinc from accessing the PLP cofactor, and any sulfur flux from NFS1 would cease, resulting in inhibition of NFS 1 activity. We showed that FXN can activate the SDU complex in the presence or absence of zinc, suggesting that FXN can cause a conformation change on ISCU to release the zinc, freeing the functional cysteine residues to begin sulfide production. This is supported by recent evidence that zinc can modulate the Fe-S cluster assembly process in the *B. subtilis* sulfur mobilization (SUF) system, whereby it stabilizes the scaffold protein SufU upon binding to cysteine desulfurase SufS.^*45*^

The zinc(II) ion observed in our ISCU samples is likely derived from recombinant expression in *E. coli*, a phenomenon that likely occurred during sample preparation for previous *in vitro* studies of recombinant *E coli* IscU and human ISCU.^*34*^, ^*37*^, ^*44*^ Some variations in the finite ratio of zinc-bound to zinc-depleted populations exist among our different preparations of recombinant ISCU, although the majority of the as-purified protein is predominantly in the zinc-bound form, which upon EDTA treatment can be converted to the zinc-depleted form resulting in modest activation of NFS 1 activity. We therefore reason that previous characterization of ISCU/IscU biochemical and biophysical properties^*34*^, ^*37*^ ^*44*^ should be interpreted with caution, and in the context that as-purified ISCU/IscU would potentially be present in both zinc-bound and zinc-depleted forms, influencing the equilibrium between ISCU conformations. Whether this also happens *in vivo*, and has a biological relevance for regulating Fe-S cluster biosynthesis by turning on/off the NFS1 reaction, is currently unknown. It is of note that in human cells, the total concentration of zinc is 200–300 *μM*,^*46*^ with the major pool of zinc residing in the mitochondria and free zinc uptake in human mitochondria reported to be in the range of 80 pm – 20 μM.^*47*^ Therefore a physiologically-relevant phenomenon could exist *in vivo*, whereby the bound zinc(II) ion could play a role in modulating Fe-S metabolism and oxidative stress in mitochondria, although this needs to be further explored.

Another objective of this study was to determine, using our recombinant expression system, the effect of substituting Met140-to-Ile (and Leu, Val, Cys) in ISCU on its biochemical and biophysical properties. The equivalent substitution in yeast Isu yielded recovery of growth and cell viability defects in cells deficient in frataxin Yfh1. Additionally, using purified yeast proteins, the IscU suppressor has been shown to activate Nfs1 and bypass the requirement for Yfh1 *in vitro*.^*42*^ Our data suggest that the human ISCU suppressor variants, when recombinantly expressed in *E. coli*, were essentially indistinguishable from the wild-type with regards to zinc(II) binding, secondary structure, thermostability, and complex formation. Importantly, under both zinc-replete and zinc-depleted buffer conditions tested, there is negligible desulfurase activity for the frataxin-free SDU complex involving either ISCU wild-type or suppressor variants.

Our inability to observe a FXN bypass effect with the human ISCU suppressor mutants may be due to differences between the yeast and human proteins or due to a number of factors, including assays and reagents, and potentially variable levels of zinc (II) that were not controlled by other studies.^*31*^–^*33*^, ^*36*^ For example, human FXN cannot activate NFS1 without ISCU present^*17*^, while yeast Yfh1 can in the absence of Isu1 (and sometimes without Isd11).^*42*^ Additionally, different activity assays were used to quantify NFS1 desulfurase activity in this study compared to the previous study reporting FXN bypass.^*42*^ Here, a colorimetric assay was used to measure sulfide production and release in the presence of a reductant, while in the previous study sulfide accumulation on NFS1 was probed with [^35^S]-Cys by scintillation counting and autoradiography. Importantly, this study has demonstrated that additional factors present in the purified samples, for example zinc(II), can directly influence NFS1 activity. Uncontrolled variability in zinc content and its impact on reported complex activities between studies, and even between batches of protein from within one study, may be an important factor that contributed to the discrepancy between studies. In summary, the complexity of the multi-component Fe-S assembly system and the level of zinc(II) co-purified with ISCU need to be taken into account in future experimental design when exploring the impact of point mutations on complex activity.

## ACKNOWLEDGEMENTS

The Structural Genomics Consortium is a registered charity (Number 1097737) that receives funds from AbbVie, Bayer Pharma AG, Boehringer Ingelheim, Canada Foundation for Innovation, Eshelman Institute for Innovation, Genome Canada, Innovative Medicines Initiative (EU/EFPIA) [ULTRA-DD grant no. 115766], Janssen, Merck & Co., Novartis Pharma AG, Ontario Ministry of Economic Development and Innovation, Pfizer, São Paulo Research Foundation-FAPESP, Takeda, and Wellcome Trust [092809/Z/10/Z]. N.G.F. and W.W.Y. are further supported by funding from the Pfizer Rare Disease Consortium. We would like to thank David Staunton at University of Oxford in the Biochemistry Department for help and data processing of the circular dichroism experiments using the DiChroweb software.

## Footnotes

* For clarity for the general scientific community, we refer to residues of human ISCU or yeast Isu following nomenclature for the description of protein sequence variants, i.e. by the numbering of the full-length protein (UNIPROT entry Q9H1K1 or Q03020). In the literature, ISCU/Isu residues has often been numbered by referring to the mature protein without the N-terminal mitochondrial targeting sequence. Both numbering schemes were included in the first mention of the residue in the main text.

